# Faster emergence behavior from ketamine/xylazine anesthesia with atipamezole versus yohimbine

**DOI:** 10.1101/337246

**Authors:** Lukas Mees, Jonathan Fidler, Matthias Kreuzer, Jieming Fu, Machelle M. Pardue, Paul S. García

**Affiliations:** Atlanta VA Center for Visual and Neurocognitive Rehabilitation, Decatur, Georgia, USA; Department of Anesthesiology, Emory University, Atlanta, Georgia, USA; Department of Anesthesiology, Technical University of Munich, Munich, Germany; Department of Biomedical Engineering, Georgia Institute of Technology, Atlanta, Georgia, USA

## Abstract

Recent interest in reversal of the hypnotic effects of anesthesia has mainly focused on overcoming a surge in GABA-mediated inhibitory signaling through activation of subcortical arousal circuits or antagonizing GABA receptors. Here we examine the reversal of anesthesia produced from non-GABA agents ketamine/xylazine and the effects of antagonists of adrenoreceptors. These antagonists vary in selectivity and produce temporally unique waking behavior post-anesthesia. We compared two antagonists with differential selectivity for α_1_-vs. α_2_-receptors, yohimbine (YOH, 1:40 selectivity) and atipamezole (ATI, 1:8500). Adult mice received intraperitoneal injections of either YOH (4.3 mg/kg), ATI (0.4 mg/kg), or saline after achieving sustained loss of righting following injection of ketamine/xylazine (ketamine: 65.0 mg/kg; xylazine: 9.9 mg/kg). Behaviors indicative of the post-anesthesia, re-animation sequence were carefully monitored and the timing of each behavior relative to anesthesia induction was compared. Both YOH and ATI hastened behaviors indicative of emergence, but ATI was faster than YOH to produce certain behaviors, including whisker movement (YOH: 21.9±1.5 min, ATI: 17.5±0.5 min, p=0.004) and return of righting reflex (RORR) (YOH: 40.6±8.8 min, ATI: 26.0±1.2 min, p<0.001). Interestingly, although YOH administration hastened early behavioral markers of emergence relative to saline (whisking), the completion of the emergence sequence (time from first marker to appearance of RORR) was delayed with YOH. We attribute this effect to antagonism of α_1_ receptors by yohimbine. Also notable was the failure of either antagonist to hasten the re-establishment of coordinated motor behavior (e.g., attempts to remove adhesive tape on the forepaw placed during anesthesia) relative to the end of emergence (RORR). In total, our work suggests that in addition to pharmacokinetic effects, re-establishment of normal waking behaviors after anesthesia involves neuronal circuits dependent on time and/or activity.

## Introduction

Reconstruction of consciousness has been studied in the context of anesthesia [1-3] and has been likened to waking from sleep [4]. Despite the active pharmacological reversal of some aspects of anesthesia such as neuromuscular blockade and opioid induced respiratory depression, the recovery of consciousness following clinical anesthesia has traditionally been considered a passive process. Recently, stimulation of arousal pathways [5] and antagonism of inhibitory signaling [6] have been investigated as potential strategies for hastening the arrival of consciousness after isoflurane anesthesia. This work has been extended to reversal of other GABA-ergic anesthetic agents such as propofol [7]. In contrast, the reversal of non-GABA agents (i.e., ketamine, xylazine, dexmedetomidine) has received much less attention, perhaps because their use as sole agents for maintenance of anesthesia is less common in human clinical practice [8-10].

Despite their lack of study, non-GABA agents remain in common use. Ketamine (K) is one of the most popular non-GABA veterinary anesthetic agents. It has been in use for over 50 years, yet there is still much to learn about its pharmacodynamic effects. Although glutamate receptors are known targets of its neurophysiologic effects (NMDA antagonism, AMPA agonism) other receptors involved in neuronal excitability have demonstrated bioactivity.

Ketamine is most often administered in combination with other anesthetics, such as xylazine, which is thought to counteract some of ketamine’s sympathomimetic effects. The intraperitoneal or intramuscular injection of xylazine in combination with ketamine is a common anesthesia technique used for procedures performed in the laboratory on mice, rats, and other animals [11-13]. Xylazine (X) specifically agonizes the α_2_-adrenoceptor [13]. Its action at the α_2_-receptor in the brain stem produces sedation through increased noradrenergic release throughout the cortex [14].

While ketamine/xylazine (K/X) is an effective combination for veterinary anesthesia, it can produce side effects such as acute hyperglycemia [15] and corneal lesions [16] if administered without a reversal agent. Currently, no pharmacologic reversal of ketamine anesthesia exists. However, in veterinary practice, the sedative actions of α_2_-agonists can be pharmacologically reversed with α_2_-antagonists, such as yohimbine and atipamezole [13]. Yohimbine (YOH) is an indole alkaloid derived from the bark of the *Pausinystaliayohimbe* tree. It has highest affinity for the α_2_-receptor, but also antagonizes the α_1_-receptor, as well as some serotonin and dopamine receptors. It is considered a selective α_2_-antagonist, with a 40:1 α_2_:α_1_ selectivity ratio. Atipamezole (ATI) is also an α_2_-antagonist, with a higher α_2_:α_1_ selectivity ratio 8500:1 [17]. ATI is also more potent than YOH. Ten times the amount of YOH is needed to block central α_2_-receptors to the same level that ATI does [18, 19].

The combination of an anesthetic cocktail and a reversal agent (of varying selectivity) can have complex influences on behavior while the animal emerges from anesthesia. A more efficient hastening of emergence with ATI compared to YOH has been suggested in the context of ketamine/xylazine anesthesia [20], but an examination of the behaviors during emergence and recovery from anesthesia with these reversal agents has yet to be studied.

Studying anesthesia produced by ketamine and α_2_-agonists has clinical relevance. Ketamine has recently experienced a renaissance in clinical usage [9] as some benefits have been demonstrated in the treatment of depression [21], attenuation of postoperative delirium [22], and reduction of opioid administration for analgesia [23]. Although xylazine is not approved for use in humans, other selective α_2_-agonists, like dexmedetomidine and clonidine, are used for blood pressure control, sedation, and as adjuncts to general anesthesia. Dexmedetomidine in combination with ketamine has been used as a preferred technique intending to minimize post-anesthesia confusion in humans [24, 25].

Here we investigated an animal model of recovery from anesthesia in the absence of GABA-ergic anesthetic drugs. We measured the appearance of behavioral markers of emergence and describe their canonical sequence of arrival following K/X anesthesia in the presence and absence of reversal by α_2_-antagonists.

## Methods

### Animals

Both male (n = 24, and female (n = 6) adult (approximately 20 – 30 grams) C57BL/6J mice (Jackson Laboratories, Bar Harbor, ME) were used. Animals were experimentally naïve and were used for only one trial each. Mice were housed under a 12:12 light:dark cycle and given standard mouse chow *ad libitum*. All procedures were approved by the Atlanta Veterans Administration Institutional Animal Care and Use Committee.

### Emergence procedure, anesthesia challenge, and early behavioral markers

Figure 1 depicts the experimental protocol. Animals were first weighed, and then placed in an open top, clear observation box for acclimation. Before induction of anesthesia, each mouse received training on a “sticky dot” test (see below). Following the training trials, mice were given an intraperitoneal injection of K/X cocktail (65 mg/kg of ketamine [Ketaved, Vedco Inc., St. Joseph, MO], 9.9 mg/kg xylazine [Anased, Akorn, Decatur, IL]), and the timer was started. Animals were considered “anesthetized” when they failed to right themselves (by placing all four paws on the surface of the chamber) after being gently placed on their back. This time was noted as the appearance of loss of righting reflex (LORR). To investigate post-anesthesia behaviors in the absence of pharmacological manipulation of alpha receptors, six mice were administered intraperitoneal ketamine (65 mg/kg) and then immediately placed into 4% isoflurane for 60 seconds which resulted in until LORR LORR (K/I regimen). Immediately after LORR, each mouse was placed on a heating pad beneath a heat lamp adjusted to maintain body temperature between 39.4-40.0 degrees Celsius. While anesthetized, adhesive tape (“sticky dot”) was applied to the right forepaw. At exactly 15 minutes after the injection of anesthesia/sedation each mouse was given either YOH [Yobine, Akorn Inc, Decatur, IL; 4.3 mg/kg], ATI [Antisedan, Orion Corporation, Espoo, Finland; 0.4 mg/kg], ATI and prazosin (ATI+PRA) [Prazosin HCl, Sigma-Aldrich, St. Louis, MO; 2.0 mg/kg], or saline (SAL) [Hospira Inc., Lake Forest, IL; 0.1mL 0.9% solution. Consequently, the groups are depicted SAL, YOH, ATI, ATI+PRA for all groups with ketamine/xylazine and K/I-SAL for the ketamine/isoflurane group in the reminder of the manuscript. The time of the following behavioral markers was taken at their first occurrence: whisker movement (any movement of whiskers), forelimb movement (any movement of either forelimb), and respiration change (either a change in rate or change in breathing depth, judged by the size of the chest excursion with each breath). Next, the time for each mouse to regain its righting reflex (RORR) was recorded. As rodents do not sleep on their backs, it is common to use RORR as an arbitrary marker for cessation of the anesthetized state (end-emergence), and so it is used here to delineate the emergence and recovery periods. Upon righting, an attempt to return the mouse to its back was performed in each mouse to ensure the righting reflex was robust. Following the RORR, each mouse was placed into a clear-walled open top box to enable observation of the recovery period behaviors.

**Figure 1.**
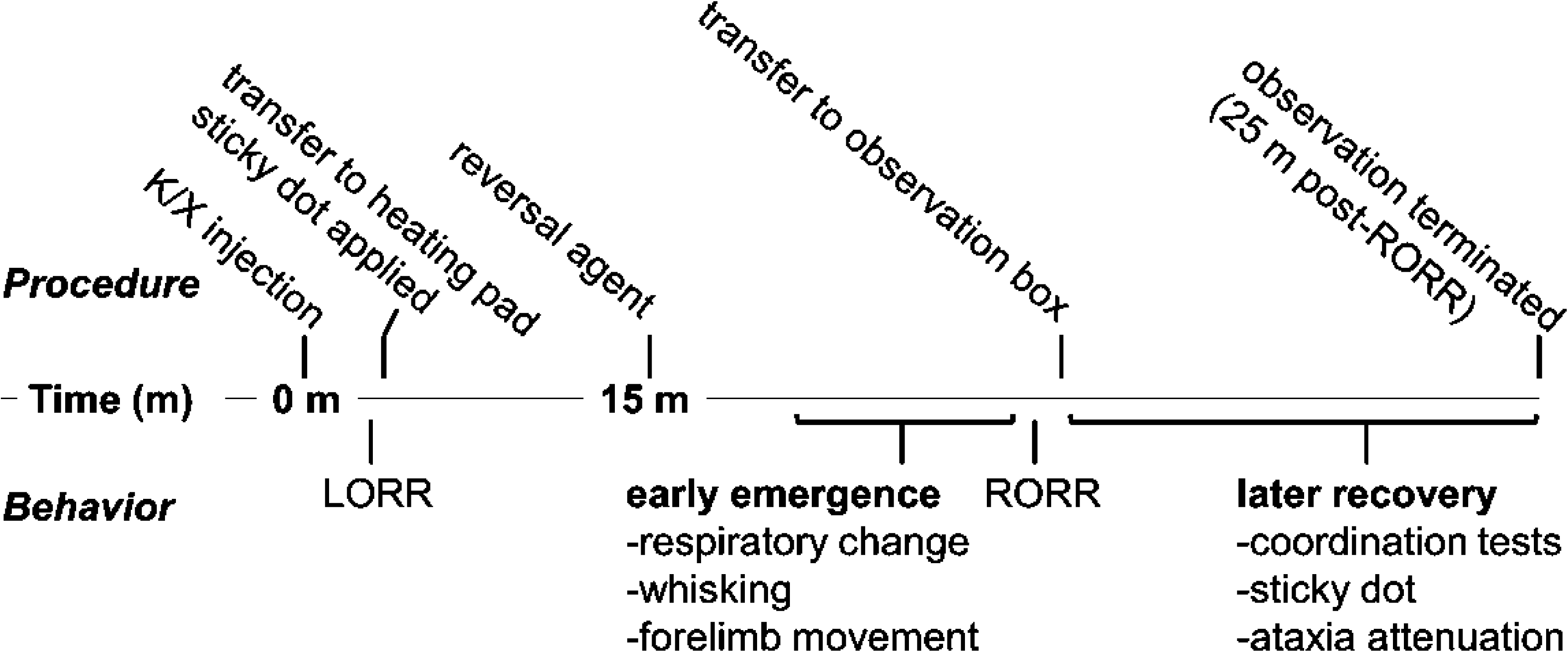
Waking behavior observation protocol. Timeline describing observation protocol. Procedures listed on top, with behaviors listed below. K/X = ketamine/xylazine, LORR = loss of righting reflex, RORR = return of righting reflex.

### Late behavioral markers and early recovery from anesthesia

The sticky dot test is a complex measure of perception and motor coordination previously used to evaluate animals after ischemic stroke [26]. Briefly, the animals receive a 2.5 × 0.5 cm adhesive tape folded over their forepaw; then the time to investigation of the tape (paw shaking or any purposeful movement towards the tape involving the nose, mouth, or alternate forepaw) is recorded. Prior to the anesthesia challenge, all mice received three trials of the sticky dot test. For our experiments we defined *recovery period* as the time between RORR and the appearance of the final marker of our observed behavioral sequence (sticky dot notice). If the animal did not attempt to remove the tape within 25 minutes after RORR, the trial ended. Ataxia was assessed at five-minute intervals after RORR by testing for splaying of the legs. This was accomplished by lifting the mouse by the tail, suspending both hind limbs and observing the hind limb reflexes after subsequent dropping of the hindquarters. Other ataxic features were recorded if present: ambulation with only the forelimbs, or otherwise uncoordinated movement. Coordinated movement was defined as diagonal cross-matched ambulation, in which the right forelimb movement was followed by movement of the left hind limb. Latency to return of diagonal cross-matched ambulation was recorded. After 25 minutes post-RORR, observation was terminated, and the mouse was sacrificed by cervical dislocation.

### Statistics

We applied non-parametric testing for evaluating our data set, because of our modest sample size. We used the Kruskal-Wallis test to evaluate possible differences between all groups and the Mann-Whitney U test as *post-hoc* test. We did not correct for multiple comparisons in order to prevent increase of false negatives [27]. But therefore, additionally to the hypothesis-based tests, we calculated the area under the receiver operating curve (AUC) together with 10000-fold bootstrapped 95% confidence intervals as effect size. As a rough estimate, according to the traditional point system, an effect can be classified as: excellent (very strong) AUC=0.9-1; good AUC=0.8-0.9; fair AUC=0.7-0.8; poor AUC=0.6-0.7; or fail: AUC=0.5-0.6. We used MATLAB R2017 (The Mathworks, Natick, MA) for our statistical tests and the MATLAB-based MES toolbox [28] to calculate AUC and 95% CI. We present our data as raw data together with the mean and the median.

## Results

### Baseline testing and induction of anesthesia

During baseline experiments of the sticky dot test all mice notice and removed the adhesive tape in less than 2 minutes by the third trial. No animal noticed or removed the tape before RORR. All 24 mice that received the K/X dose described in the methods experienced LORR in less than 5 minutes.

### **Emergence from ketamine/xylazine anesthesia is hastened with the administration of α_2_**-**antagonists**

Among the early behaviors observed before RORR, change in respiration rate, whisker movement, and forelimb movement were recorded. Figure 2 (A-C) contains the detailed information. ATI and YOH produced signs of waking earlier than SAL in all three behaviors. The time to whisker movement was different among the groups (p=0.0005, χ^2^=15.16). Compared to saline (n=6) treated animals, YOH(n=6, p=0.0022, 22.2 [19.3 23.7] min) and ATI (n=6, p=0.0022, 17.4 [17.0 18.3] min) showed faster recovery to whisker movement than SAL (46.8 [41.0 55.3] min).AUC indicated perfect separation (very strong effect) between the groups, i.e., AUC=1. We observed the same result when comparing the ATI with the YOH group (p =0.0022; AUC=1, very strong effect). Time required for anesthetized mice to exhibit a change in respiration was also different among groups (p=0.0013, χ^2^=13.35) depending on reversal agent. We derived very similar statistical results for time to respiration as for time to whisker movement. The SAL group took significantly longer to express respiration signs (n=6, 47.1 [41.0-55.3] (median [min max] min) when compared to YOH (n=5, 22.2 [19.2-24.5] min) with p=0.0043 and when compared to ATI (n=5, 17.5 [17.0-18.3] min with p=0.0043. AUC also showed perfect separation (AUC=1) between the groups. When compared to YOH, ATI animals reached this behavioral milestone significantly earlier as well (p=0.0079, AUC=1).Overall, ATI and YOH have similar profiles in the sequence of early markers of emergence.

**Figure 2.**
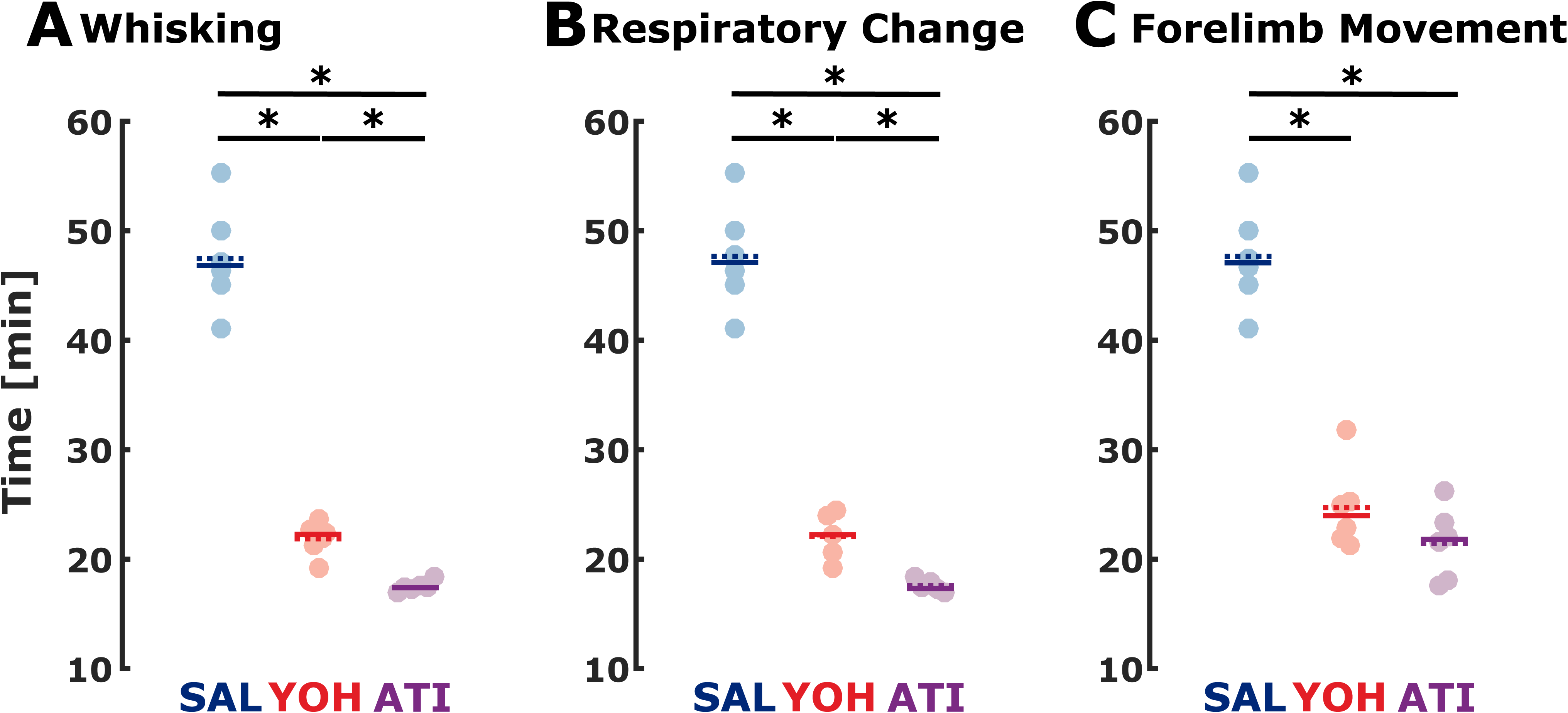
Both YOH and ATI reduce the time required to exhibit the first behavioral signs of emergence from ketamine/xylazine anesthesia. Time for first incidence of individual waking behaviors is plotted, including whisker movement (A), respiratory change (B), and forelimb movement (C). Time measurements are from ketamine/xylazine injection. * indicate significance between different groups; solid bar: median, dashed bar: mean.

### Yohimbine increases the time to completion of the emergence sequence

The timing of RORR was significantly different among treatments (p=0.0011; χ^2^=13.66) as shown in Figure 3. The SAL group took longer to right themselves after ketamine/xylazine anesthesia (56.4 [46.0-63.2] min) compared to YOH (38.7 [29.8-51.2] min) with p=0.0260 and AUC=0.89 [0.67 1] (strong effect) and ATI (25.9 [24.6-27.4]) with p=0.0022 and AUC=1 (very strong effect). ATI animals also exhibited RORR faster than YOH animals (p=0.0022 and AUC=1 (very strong effect). Interestingly, the time delay between the first exhibited behavior and RORR showed significant difference among groups (p=0.0248; χ^2^=7.40), but with a different pattern. YOH showed a significantly longer delay in completion of the emergence sequence (n=6, 17.1 [7.9-29.3] min) compared to SAL (n=6, 8.8 [1.0-13.6] min) with p=0.0152 and AUC=0.92 [0.67 1] and ATI (n=6, 8.1 [7.2-10.3] min) with p=0.0260 and AUC=0.89 [0.58 1]. There was no significant difference between SAL and ATI (p=1; AUC=0.5 [0.17 0.83]). This suggests that although yohimbine hastened the start of emergence behaviors, completion of the entire sequence of emergence behaviors was lengthened by administration of yohimbine (Figure 3B).

**Figure 3.**
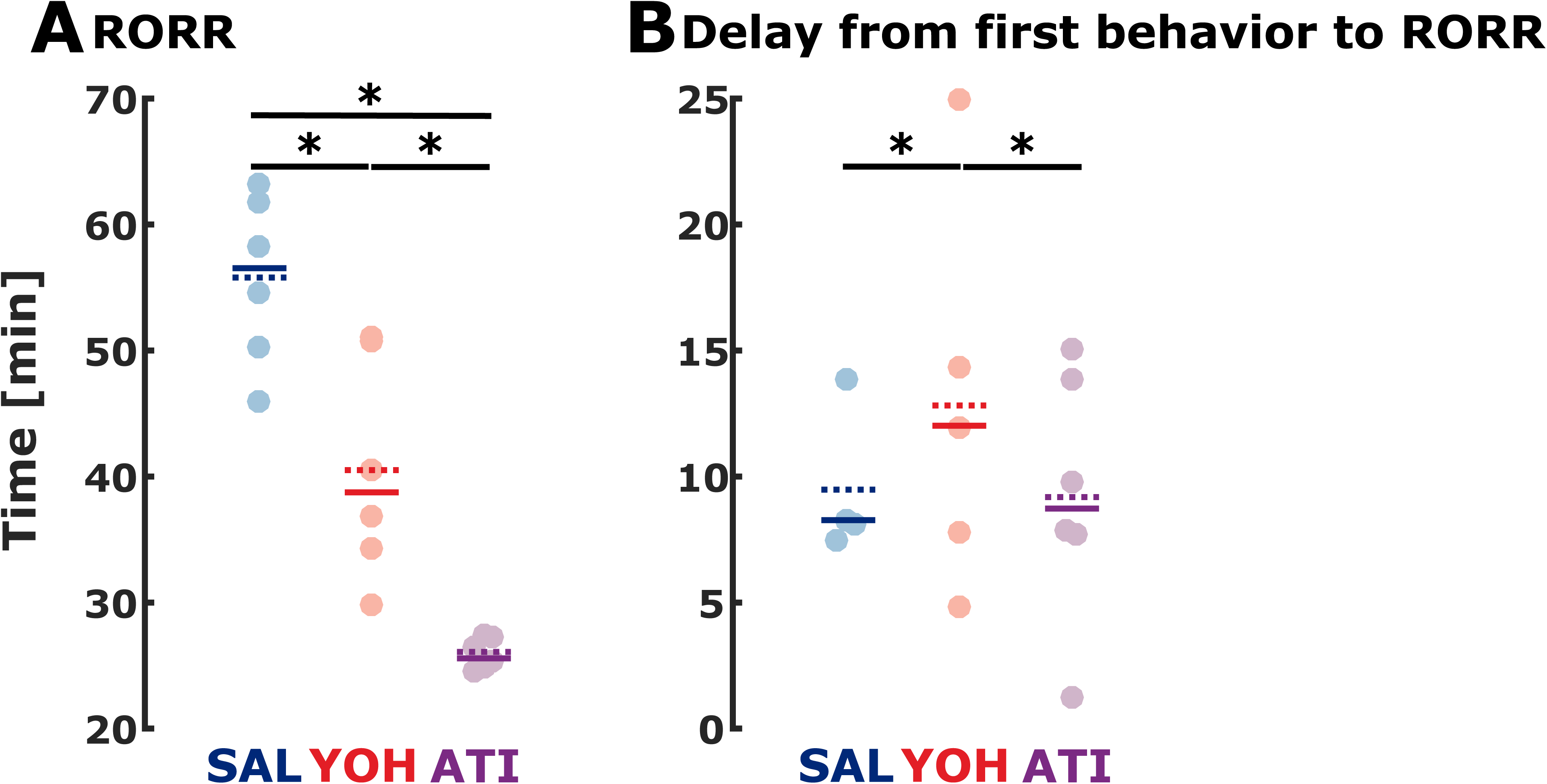
Both α2 antagonist treated groups recover righting reflex faster, but YOH has increased delay between early markers and RORR. (A) Return of righting reflex. (B) Delay from first behavioral marker to RORR. Time measurement in (A) is from ketamine/xylazine injection and from the first exhibited behavior in (B). * indicate significance between different groups; solid bar: median.

### Atipamazole and yohimbine hasten recovery from ketamine/xylazine anesthesia

The recovery of locomotor activity in an uncoordinated fashion (uncoordinated movement) was significantly different among treatment groups (p=0.0008, χ^2^=14.36). YOH mice exhibited uncoordinated movement faster (n=6, 41.3 [34.3-51.2] min) than SAL mice (n=6, 59.6 [49.7-65.7] min) with p=0.0087 and AUC =0.94 [0.78 1] (very strong effect), while ATI mice showed uncoordinated movement earlier (n=6, 28.6 [25.4-32.4] min) than both SAL (p=0.0022; AUC=1 (very strong effect)) and YOH (p=0.0022; AUC=1 (very strong effect); Figure 4A). Latency to the first notice of the sticky dot was different between groups (p=0.0057, χ^2^=10.32). ATI mice were faster (n=6, 36.1 [26.1-40.4] min) to identify the sticky dot compared to both SAL (n=4, 67.1 [54.1 70.0] min) with p=0.0095 and AUC=1 (very strong effect) as well as and YOH (n=6; 59.0 [39.2-62.7] min) with 0.0087 and AUC=0.97 [0.8 1] (very strong effect). YOH mice were statistically indistinguishable from SAL mice in noticing the sticky dot (p=0.19, AUC=0.8 [0.4 1] (strong, but not significant effect); Figure4B). Two saline-treated mice failed to show diagonally cross-matched ambulation, notice the sticky dot, or show ataxia attenuation within 25 minutes after RORR and one yohimbine-treated mouse did not notice the sticky dot within 25 minutes after RORR. These three animals were removed from the analysis in Figure 4B (see also supplemental Figures). But for complete presentation of the results without removed animals, we set the times of appearance of behavioral markers these animals to the maximum RORR+25 min. The results were similar to the reduced data set. The time to sticky dot notice was significantly different among the groups (p=0.0016, χ^2^=12.88). Time to event was 69.2 [54.1 88.2] min for SAL, 57.0 [39.2 62.7] min for YOH, and 36.1 [26.1 40.4] min for ATI. The pairwise comparison led to p=0.0411, AUC=0.86 [0.58 1] for SAL vs. YOH, to p=0.0022, AUC=1 for SAL vs. ATI and to p=0.0043, AUC=0.97 [0.83 1] for YOH vs. ATI. The recovery of more coordinated (diagonally cross-matched ambulation) locomotor efforts was different between groups (p=0.0005, χ^2^=10.65). YOH mice (n=5, 49.1 [39.3-54.3] min) showed a trend towards faster diagonally cross-matched ambulation than SAL controls (n=4, 66.7 [51.0-71.8] min) with p=0.0635 and AUC =0.9 [0.6 1] (very strong effect), while ATI mice (n=6, 36.4 [30.4-37.4] min) were faster than SAL (p=0.0159, AUC=1, very strong effect) and YOH mice (p=0.0080, AUC=1, very strong effect); Figure S1). The time delay to ataxia attenuation was similarly dependent on treatment (p=0.0043, χ^2^=10.88). YOH mice exhibited ataxia attenuation earlier (n=6, 47.7 [39.3-56.2] min) than SAL mice (n=4, 66.4 [51.0-71.7] min) with p=0.0381 and AUC=0.92 [0.67 1] (very strong effect), while ATI mice (n=6, 36.8 [29.6-42.4] min) were faster to show ataxia attenuation compared to both SAL p=0.0095 and AUC=1 (very strong effect) and YOH (p=0.0152, AUC=0.92 [0.70 1], very strong effect; Figure S1).

**Figure 4.**
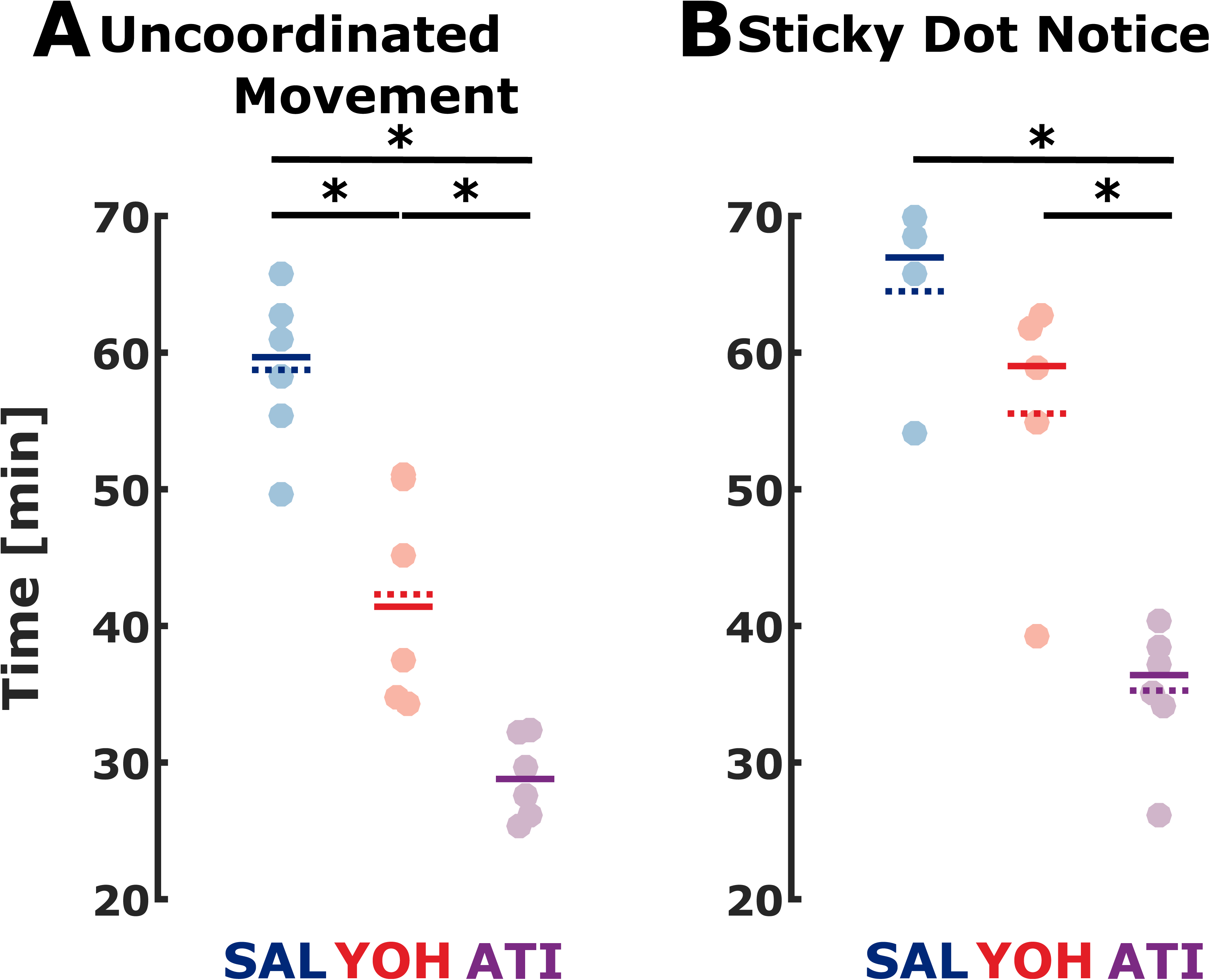
Appearance of motoric behaviors during recovery show ATI elicits activity faster than both YOH and SAL. Time delay to post-RORR behaviors are compared, including uncoordinated movement (A) and notice of sticky dot (B). Time measurements are from ketamine/xylazine injection. * indicate significance between different groups; solid bar: median.

### Emergence andrecovery behaviors are influenced by activity at α_1_-receptors

Three animals (2, SAL and 1, YOH) did not regard the sticky dot before the experiment timed out (25 minutes after RORR) so a maximum of 25 minutes was assigned as their recovery period. We did not find a significant different distribution in times from RORR to sticky dot notice between the groups (p=0.5273, χ^2^=1.28). Figure 5 graphs the latencies which were 11.1 [7.5-25] min for SAL (n=6), 13.2 [4.9-25.0] min for YOH (n=6), and 8.8 [1.2-15.0] min for ATI (n=6). The pairwise *post hoc* comparisons did not reveal any significant changes between the groups, neither did the AUC analysis reveal any trends. As an additional piece of information, the latencies between RORR to uncoordinated movement, diagonally cross matched ambulation and ataxia attenuation can be found in Figure S2. The observed difference between ATI and YOH during emergence (Figure 3B) and the observation that no animal timed out during recovery for ATI, but one did for YOH (Figure 5) prompted us to further examine the role of α1-receptors in these post-anesthesia behaviors.

**Figure 5.**
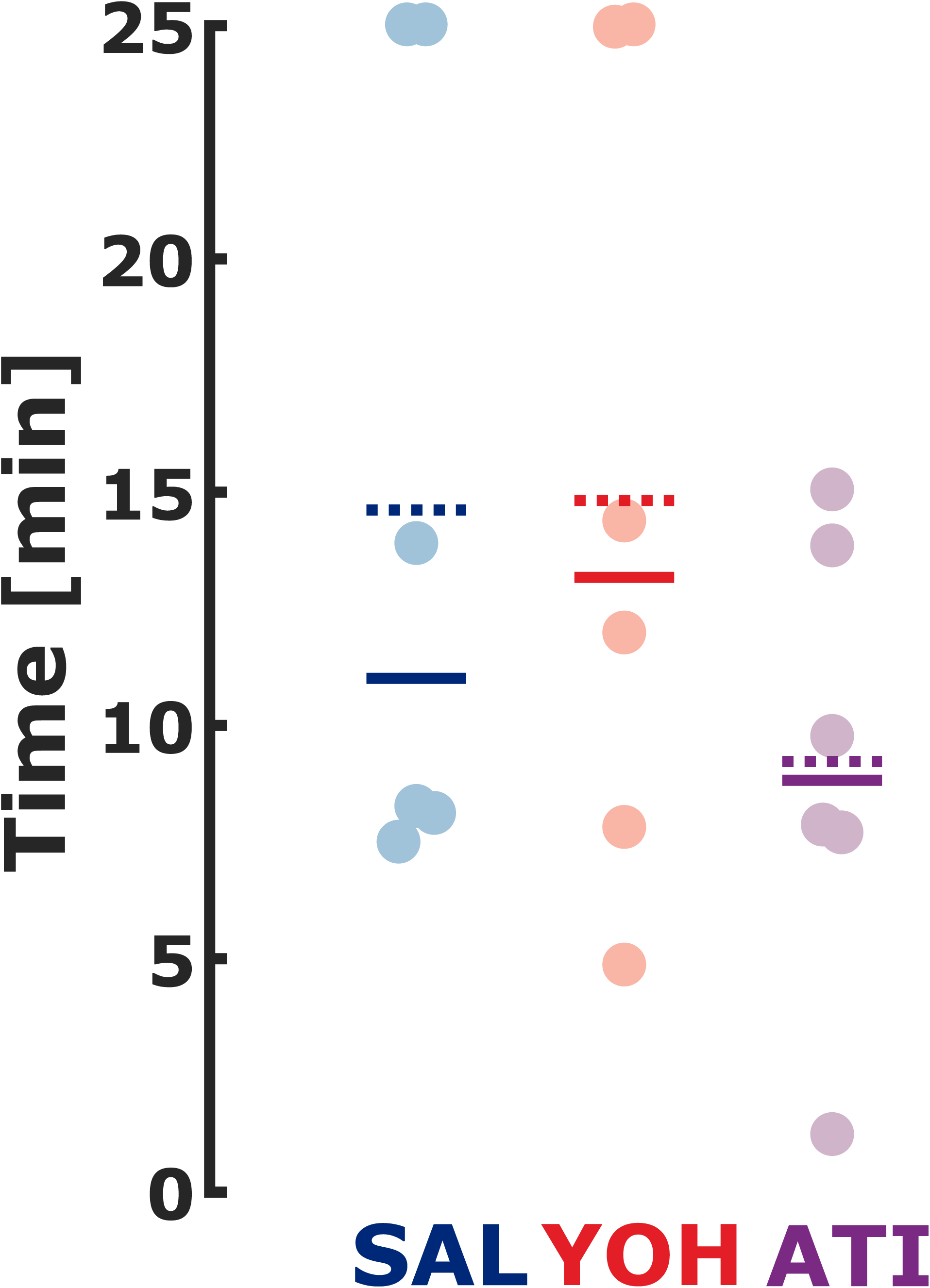
Time from RORR to sticky dot notice (recovery period) between treatment groups. Latency from RORR to the first notice of the sticky dot. * indicate significance between different groups; solid bar: median.

### Prazosin, a selective α1 inverse agonist does not hasten emergence from ketamine/xylazine anesthesia in the presence of atipamezole

Co-administration of prazosin (2.0 mg/kg) with atipamezole does not hasten RORR but prolongs sticky dot removal in the recovery from ketamine/xylazine anesthesia. Time to RORR for ATI+PRA was 26.1 [18.2 52.1] min and hence not significantly different from the ATI group (25.9 [24.6 27.4] min: p=0.7922; AUC=0.57 [0.20 0.93]; Fig. 6A). The time to sticky dot notice for ATI+PRA was 60.8 [33.3 81.7] min and also not significantly different from the ATI group (36.1 [26.1 40.4] min; p=0.2468; AUC=0.73 [0.33 1] (fair effect); Fig. 6B). However, the AUC>0.7 may indicate a trend towards a longer time to sticky dot notice with ATI+PRA. The time from the first marker of emergence to RORR was 9.3 [1.3 34.9] minfor the ATI+PRA group and hence not significantly different from the ATI group (8.1 [7.2 10.3] min; p=0.8918; AUC=0.53 [0.13 0.90], Fig. 6C). The recovery period, i.e., the time from RORR to sticky dot notice, was 25.4 [13.2 54.5] min for the ATI+PRA group. This was significantly slower (8.8 [1.2 15.0] min; p=0.0173; AUC=0.93 [0.73 1] (very strong effect)) compared to the ATI group (Fig. 6D).

**Figure 6.**
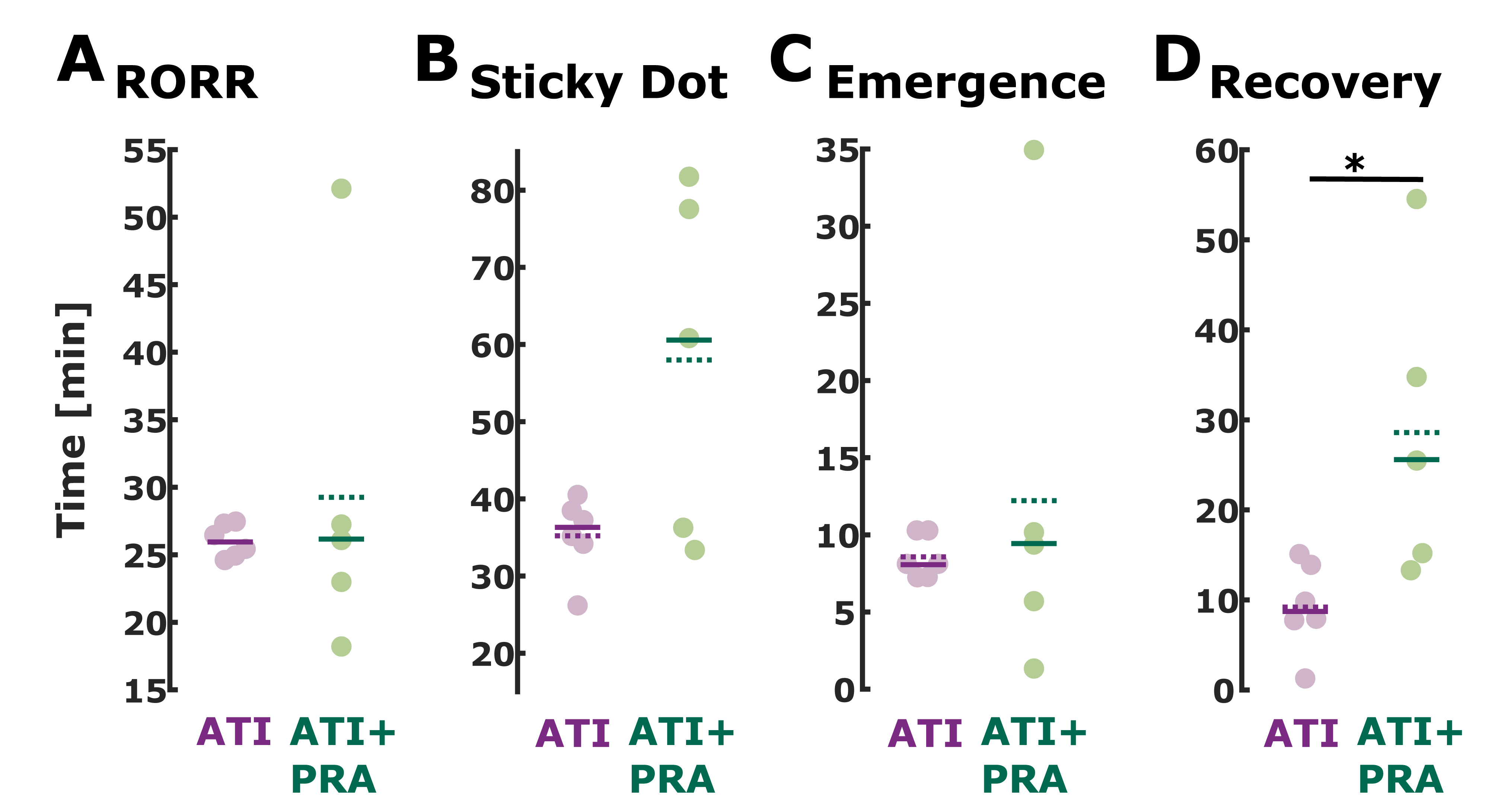
Latency to RORR and sticky dot notice and the emergence and recovery period duration for the ATI and ATI+PRA group. There were no significant differences between the ATI and ATI+PRA group in the latency to RORR and sticky dot notice. There was no difference in the duration of the emergence period as well. The recovery period was significantly shorter for ATI than for ATI+PRA.

Through a disruption of normal α_1_-receptor activity, the recovery from ketamine/xylazine in the ATI+PRA group is lengthened. In similar fashion, the emergence period (whisking to RORR) is increased for animals given YOH for reversal (Figures 6C, 6D). This highlights the importance of alpha receptor pharmacology in emergence and recovery from this ketamine/xylazine regimen.

### Lengthy recovery is associated with effects on α1 receptors

These mice were given a brief exposure to isoflurane to induce LORR and an equivalent amount of ketamine to the K/X regimen. Figure 7 compares the latency to noticing the sticky dot for the ketamine/isoflurane regimen with sham (saline) reversal (K/I-SAL). The Kruskal-Wallis test indicated a significant difference among groups (p=0.0052, χ2: 14.79). Time to sticky dot notice for KET was significantly shorter when compared to SAL (p=0.010, AUC=1, very strong effect), YOH (p=0.009, AUC=0.97 [0.80 1], very strong effect), and ATI+PRA (p=0.0303, AUC=0.9 [0.67 1], strong to very strong effect). There was no significant difference when compared to the ATI group (p=0.180, AUC=0.75 [0.42 1], fair effect). This is the group that was given only α_2_-selective agonists and antagonists and like the K/I-SAL regimen no α_1_-antagonism. Figure 8 is a summary of all the emergence and recovery observations for all 5 regimens. Qualitatively, the appearance of emergence and recovery behaviors indicative of a return to neurocognitive baseline varied in time but the order of these behaviors was largely unaltered across groups. To complete the picture, Figure S3 presents a model of the pharmacodynamic effects and the latencies of emergence and recovery period for all groups.

**Figure 7.**
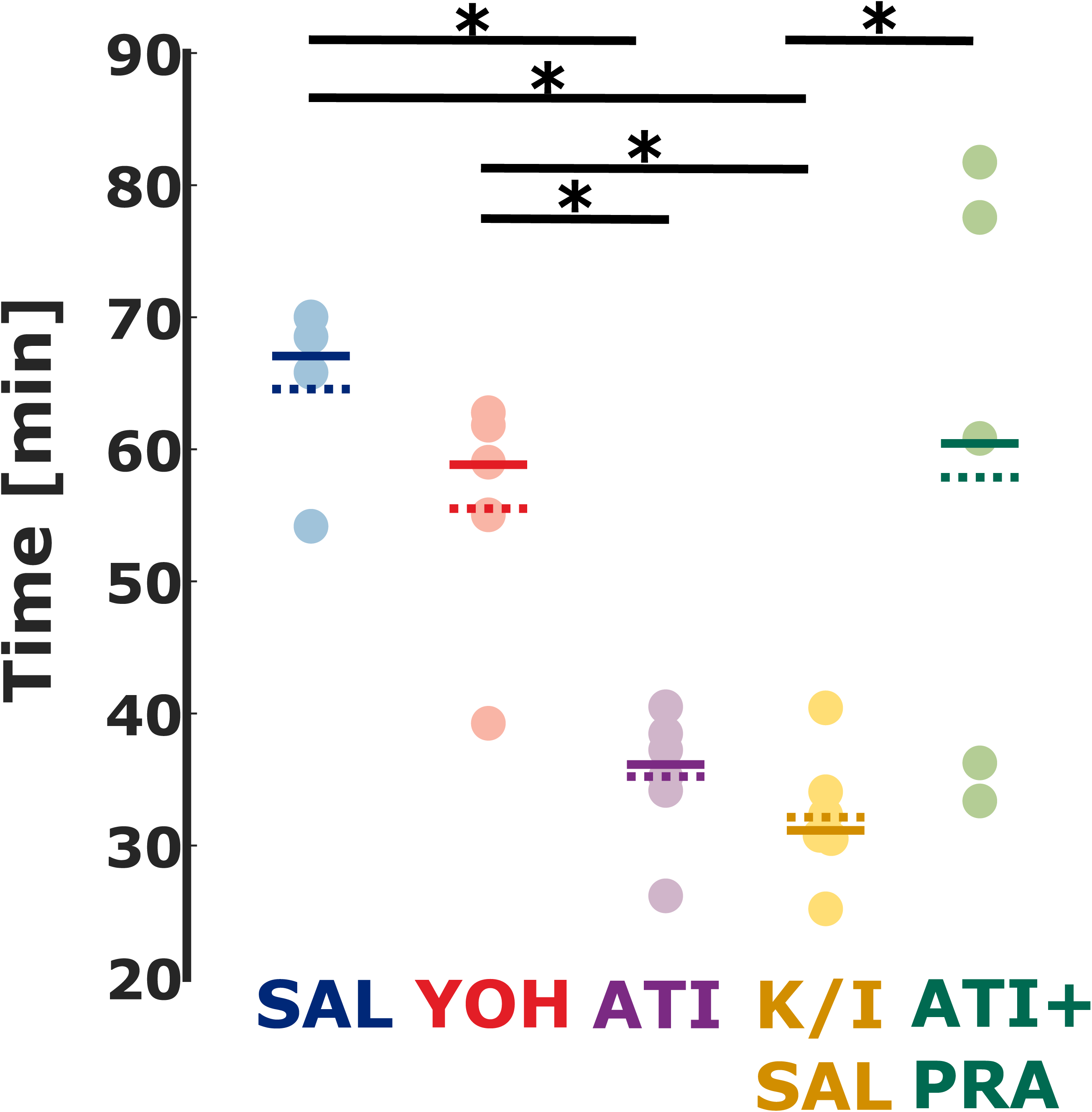
Timing of waking behavior incidence varies with reversing agent, while order is generally maintained. Animals that did not receive alpha receptor agonists or antagonists (KET+ISO+SAL) notice the sticky dot earlier than animals from the SAL and YOH group The Kruskal-Wallis test indicated a significant difference among groups (p=0.0052, χ2: 14.79). The significant differences between the groups exclusive K/I-SAL are reported in the results section. Time to sticky dot notice for K/I-SAL was significantly shorter when compared to SAL (p=0.010, AUC=1), YOH (p=0.009, AUC=0.97 [0.80 1]). There was no significant difference when compared to the ATI group (p=0.180, AUC=0.75 [0.42 1])

**Figure 8.**
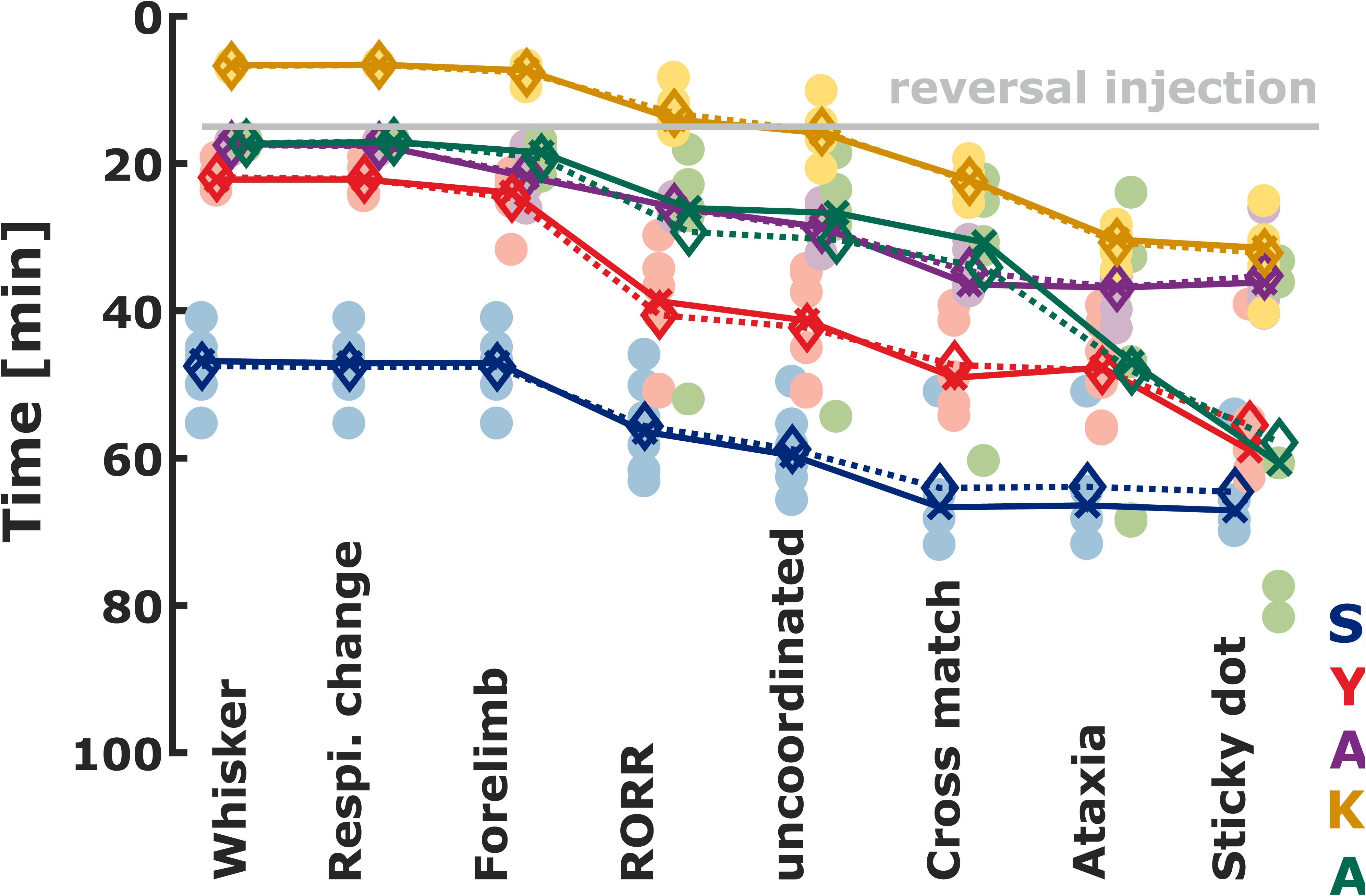
Timing of waking behavior incidence varies with reversing agent, while order is generally maintained. All measured waking behavior hallmarks compared between mice receiving SAL (blue), YOH (red), or ATI (purple). The gray line indicates time of reversal injection, 15 minutes post-ketamine/xylazine injection. Solid lines connect the medians and dotted lines connect the means.

## Discussion

In this study, behavioral milestones indicative of the approach to normal neurocognitive function after anesthesia with ketamine/xylazine were observed following injection of reversal agents. As predicted, both YOH and ATI effectively shortened the time required to reach these milestones during emergence from anesthesia compared to saline.

The behavioral profile between YOH and ATI mice during emergence and recovery from anesthesia was not identical, in agreement with previous studies [20]. Differential pharmacology between YOH and ATI offers some possible explanations for this.First, there are slight differences between YOH and ATI affinity for α_2_-receptor subtypes. ATI has an equal affinity for α_2a,_α_2b,_α_2c,_ and α_2d_, while YOH has similar affinity for all the α_2_-subtypes except for α_2d_, for which it has a lower affinity. Xylazine indiscriminately targets all of the α_2_-subtypes [29]. Due to YOH’s lower affinity for the α_2d_-subtype, it is possible that some α_2_-receptors could still be agonized by xylazine resulting in a prolonged emergence as compared to ATI.

While both drugs predominately target α_2_-receptors, YOH interacts with other systems. Unlike ATI, which has a negligible affinity for serotonin (5-HT) [17, 18], β_1_/β_2_-adrenergic, muscarinic, dopamine_2,_ tryptamine, GABA, opiate or benzodiazepine receptors [17], YOH is less discriminatory. High doses of YOH (>1 mg/kg; the current study used 4.3 mg/kg) have been shown to have 5-HT1A agonistic properties, which lead to decreases in heart rate, blood pressure, activity level, and body temperature [30]. Similarly, in doses approximating those used in the current study, YOH has been shown to decrease ambulation, an effect not seen with more selective α_2_-antagonists [19]. During the post-RORR period in the observation box, YOH and ATI animals exhibited qualitatively different levels of exploratory behavior.

In an attempt to characterize non-α_2_-interactions, the current study dosed the reversal agents in order to equalize α_2_blockade between the two reversal agents. Pertovaara et al. found that,in order to block α_2_ receptors to the same level that ATI does, about ten times the amount of YOH needs to be administered [18, 19]. We dosed 10.8 times more YOH compared to ATI. This provides some evidence that non-α_2_-interactions in YOH are slowing down the emergence process, while possibly leading to hypoactivity during recovery.

Because YOH has an α_2_:α_1_ selectivity ratio over 200 times smaller than ATI [17], α_1_ antagonism could be implicated in YOH mice. Antagonism of α_1_-receptors is mechanistically involved in the sedating effects of some anti-psychotics (quetiapine, risperidone) [31] as well as anti-hypertensives. Some of the behavioral effects of co-administering the selective α_1_-inverse agonist prazosin [32] with atipamezole mimicked the yohimbine reversal, specifically the time to notice of sticky dot, the final behavioral marker we observed. This suggests that the slowing of recoveryafter ketamine/xylazine anesthesia after yohimbine reversal (relative to ATI) could be mediated by antagonism of α_1_-receptors and likely not subsequent hypoactivity due to the effect of yohimbine on 5-HT, or other receptors. Our experiments with ketamine in the absence of xylazine or any other adrenoreceptor manipulation further support the notion that α_1_-receptor antagonism can delay complex behaviors indicative of the restoration of normal behaviors post-anesthesia (notice of sticky dot). In situ hybridization experiments reveal that neurons in layers II-V of most areas of the cerebral cortex the lateral amygdala, hippocampus, and reticular thalamus all have high density of α_1_ receptors [33]. Distribution of normal signaling of these regions may contribute to failure to achieve a normal recovery sequence efficiently.

While ATI mice were more active than YOH mice, they showed a profound lack of coordination once righted. This ataxia did not completely resolve during the 25-minute observation. α_2_-adrenoceptors are known to be present in the cerebellum [34]. Further studies will be necessary to determine if this prolonged ataxia is related to the hastening of emergence, causing enhanced locomotion before coordination is re-established, or a lingering effect of atipamezole on cerebellar function.

A limitation of our study is the failure to pharmacologically antagonize the effects of ketamine. Although ketamine is known to inhibit NMDA receptors, it is pharmacologically promiscuous and its exact mechanism for producing surgical anesthesia is unknown. Unlike xylazine, dexmedetomidine, opioids, and benzodiazepines, no pharmacologic agent specifically reverses all of thepharmacodynamic effects of ketamine. Some evidence suggests that ketamine minimally interacts with adrenoceptors [35], but these interactions have yet to be thoroughly examined. While it is not possible from this data to distinguish the effects of residual ketamine from lingering effects of the antagonists, our observations of mild hyperactive ataxic behavior in ATI animals are similar to the clinical situation often described for recovery from ketamine anesthesia characterized by excitation and features of emergence delirium [36]. This supports the notion that ATI treated animals may be exhibiting behavior typical of ketamine after effectively eliminating xylazine’s pharmacodynamic effects (Figure S3). Interestingly, ataxia appeared to be attenuated after approximately the same delay (in reference to RORR as opposed to anesthesia injection) in all groups. In parallel, if measured in reference to RORR, other measures of coordination recovery (uncoordinated movement, diagonally cross-matched ambulation) and higher perception and motor processing (sticky dot notice) had the same delay across groups. It appears that, although ATI produces a more efficient emergence from ketamine/xylazine anesthesia, it does not improve late recovery compared to YOH or even no reversal agent at all (Figure 8). It is possible that the differences in waking behaviors between ATI and YOH are arising from a differential clearance in ketamine and xylazine, rather than off-target interactions. It would be excellent to characterize these effects further with additional experiments examining the potential for a dose-effect of YOH and/or ATI. Based on our findings we conclude that proper reconstruction of network activity, requiringspecific activation of networks involving adrenoreceptors, underlies restoration of coordinated movement, as opposed to this being solely a consequence of ketamine pharmacokinetics. Future studies involving careful blood sampling over time would be necessary to determine which has the greatest influence.

Although both YOH and ATI produced waking behaviors before saline, ATI was slightly quicker to elicit several markers. These differences are likely attributable to the differential affinity between the two drugs for α_2_-subtypes, as well as α_1_-interactions. Because theeffects of adrenoceptor antagonism on behavior are dose-dependent [19, 30], further experiments are needed to compare these results to lower doses of these drugs. During recovery from anesthesia it is difficult to determine if the animal is attempting to explore their environment versus exhibiting an escape response, however motoric behaviors can still be observed and measured. Quantification of arousal, exploratory behavior, and balance should be done given the observations made during the current experiment. In total, our results highlight that an efficient emergence is not necessarily a preferred trajectory for the immediate post-anesthesia recovery. In addition to pharmacokinetic effects, the re-establishment of normal behaviors after anesthesia likely involves neuronal circuits dependent on time and/or activity.

## Figure Legends

**Figure S1. Relative to ketamine/xylazine injection, ATI hastens diagonally cross-matched ambulation and ataxia attenuation faster than both SAL and YOH.** A) latency to diagonally cross-matched ambulation; B) latency to ataxia attenuation. Time measurements are from ketamine/xylazine injection. Individual animals are plotted as translucent shapes overlaying lines representing the mean (dashed) and the median (solid). * indicate significance between different groups; the (*) indicates a non-significant p=0.063 (MWU), but a very strong effect as indicated by an AUC=0.90 [0.60 1].

**Figure S2. Time from RORR to sticky dot notice (recovery period) shows no difference between treatment groups.** Latency from RORR to the first notice of the sticky dot. * indicate significance between different groups; solid bar: median

**Figure S3. YOH lengthens time to complete emergence and recovery.** The upper graph is an idealized model that schematically depicts the expected pharmacodynamic effects (estimated overall effect of the drugs on the animal) of anesthetic agents over time (x-axis on the same scale as lower graph). Ketamine = yellow line, xylazine = green, xylazine with reversal agent = light green line. Mean latency for emergence period (whisker movement to RORR, solid lines) and recovery period (RORR to sticky dot notice, dashed lines) is plotted on the lower graph. SAL = blue, YOH = red, ATI = purple, K/I-SAL = yellow, ATI+PRA = green. Mean values are plotted. Vertical dashed gray line represents time of reversal agent injection.

